# Conditional degradation of UNC-31/CAPS enables spatiotemporal analysis of neuropeptide function

**DOI:** 10.1101/2022.07.14.499970

**Authors:** Rebecca Cornell, Wei Cao, Jie Liu, Roger Pocock

## Abstract

Neuropeptide release from dense-core vesicles in *Caenorhabditis elegans* is promoted by UNC-31, ortholog of the calcium-dependent activator protein for secretion (CAPS). Loss of UNC-31 causes multiple phenotypes in *C. elegans* including reduced motility, retention of late-stage eggs and reduction in evoked synaptic release. However, the ability to analyze UNC-31 function over discrete timescales and in specific neurons is lacking. Here, we generated and validated a tool to enable UNC-31 expression and spatiotemporal functional analysis. We show that endogenously tagged UNC-31 is expressed in major ganglia and nerve cords from late-embryonic stages through to adult. Using the auxin-inducible degradation system, we depleted UNC-31 post-embryonically from the nervous system and revealed defects in egg-laying, locomotion and vesicle release that were comparable to *unc-31* null mutant animals. In addition, we found that depleting UNC-31 specifically from the BAG sensory neurons causes increased intestinal fat storage, highlighting the spatial sensitivity of this system. Together, this protein degradation tool may be used to facilitate studies of neuropeptide function at precise cellular and temporal scales.

**Significance statement:** Animal behavior and physiology is controlled by neuropeptides that are released from specific neuronal sources. The ability to dissect discrete neuropeptide functions requires precise manipulation of neuropeptide release. We have developed and validated a tool that enables precise spatiotemporal regulation of neuropeptide release that will enable researchers to examine neuropeptide function at exceptional resolution.

## Introduction

Neuronal stimulation causes the release of fast-acting neurotransmitters and slower-acting neuropeptides. Calcium (Ca^2+^) entry into neurons precedes neurotransmitter and neuropeptide exocytosis, with neurotransmitters released from synaptic vesicles (SVs) and neuropeptides from dense-core vesicles (DCVs). UNC-31, the *C. elegans* ortholog of calcium-dependent activator protein for secretion (CAPS), is a phosphoinositide-binding protein that is a critical for neuropeptide release from DCVs (Charlie et al., 2006; Gracheva et al., 2007; Sieburth et al., 2007; Speese et al., 2007). UNC-31 houses conserved C2 (coiled coil) and PH (pleckstrin homology) domains that coordinate tethering to the plasma membrane, a MUN (Munc13-homology) domain that interacts with syntaxin across phyla and a dense-core vesicle binding domain (DCVBD) (Speese et al., 2007). Loss of UNC-31 causes synaptic DCV accumulation, reduced neuropeptide release, and defective behaviors associated with reduced neuropeptide signaling such as uncoordinated locomotion and retention of late-stage eggs (Charlie et al., 2006; Gracheva et al., 2007; Sieburth et al., 2007; Speese et al., 2007). In addition, *unc-31* null mutant animals exhibit impaired evoked SV release that is likely an indirect effect of reduced neuropeptide action via the Gα_s_ pathway (Gracheva et al., 2007). Genetic screens have previously generated multiple *unc-31* mutant alleles to enable domain-specific functional analysis (Speese et al., 2007). However, the availability of tools to enable the manipulation of UNC-31 expression in a spatiotemporal manner is lacking. We endeavored to generate a genetic tool to enable 1) visualization of UNC-31 endogenous expression and localization and 2) temporal and spatial depletion of endogenous UNC-31.

The auxin-inducible degradation system in *C. elegans* is a powerful method that enables precise spatiotemporal manipulation of gene expression to facilitate functional studies (Zhang et al., 2015; Cao et al., 2021). The system leverages the ability of the *Arabidopsis thaliana* auxin hormone to regulate protein expression by activating the F-box transport inhibitor response 1 (TIR1) protein - the substrate recognition component of a Skp1-Cullin-F-box E3 ubiquitin ligase complex - which ubiquitinates degron-tagged proteins for proteasomal degradation (Ruegger et al., 1998; Gray et al., 1999; Dharmasiri et al., 2005). In *C. elegans*, the system requires the target protein to be endogenously tagged with the plant auxin-inducible degron (AID) sequence, expression of the plant TIR1 F-box protein in the tissue/cell of interest and incubation with auxin (Zhang et al., 2015). Conditional depletion of *C. elegans* proteins can be precisely controlled by the spatial expression of TIR1, timing of auxin application, and degron tagging of specific protein isoforms (Zhang et al., 2015). In addition, depleted proteins can recover their expression when animals are removed from auxin. However, the rate at which this occurs can depend on the transcriptional/translation rates of the gene/protein of interest, the concentration of auxin used, and the developmental stage of treatment.

Here, we describe the generation and validation of a tool to enable UNC-31 conditional depletion. Endogenous tagging of UNC-31 with GFP-degron-TEV-3xFLAG reveals that expression commences during late embryogenesis and continues throughout adulthood. As reported previously using UNC-31 immunofluorescence (Gracheva et al., 2007), correct localization of endogenously-tagged UNC-31 protein is dependent on the *C. elegans* kinesin UNC-104. We further show that addition of the GFP-degron-TEV-3xFLAG tag does not cause detectable phenotypes associated with *unc-31* loss. We demonstrate the effectiveness of this conditional UNC-31 allele by analyzing the kinetics of auxin-induced UNC-31 depletion and revealing that post-embryonic UNC-31 depletion phenocopies *unc-31* null mutant phenotypes (egg-laying, locomotion, and evoked excitatory postsynaptic currents (EPSCs)). In addition, we show that depletion of UNC-31 from individual neurons (BAG sensory neurons) can reveal neuron-specific neuropeptide functions (intestinal fat storage). Together, we provide a tool that enables inhibition of neuropeptide release in a spatial and temporal manner, which will help accelerate functional dissection of neuropeptide functions.

## Materials and methods

### Genetics

*C. elegans* strains were cultured on Nematode Growth medium (NGM) plates and fed with *Escherichia coli* OP50 bacteria at 20°C, unless otherwise stated (Brenner, 1974; Avery et al., 1993; Liu et al., 2009). Strains used: Bristol N2 (wild-type), RJP5269 *rp166(unc-31-linker-GFP-TEV-AID-FLAG) IV, RJP5287 rp166 IV; him-5(e1490) V, RJP5296 reSi7(Prgef-1::TIR1::F2A::mTagBFP2::NLS::AID::tbb-2 3’UTR) I; rp166 IV, EG5096 oxIs364(unc-17p::channelrhodopsin::mCherry) X, RJP5382 reSi7(Prgef-1::TIR1::F2A::mTagBFP2::NLS::AID::tbb-2 3’UTR) I; rp166 IV; oxIs364 X, CB928 unc-31(e928) IV, RJP5601 unc-104(e1265) II; rp166 IV*.

### CRISPR-mediated generation the *gfp-tev-degron-flag-unc-31* strain

An N-terminal GFP-degron-FLAG-UNC-31 knock-in strain (*rp166)* was generated using CRISPR/Cas9-triggered homologous recombination (Dokshin et al., 2018). The crRNA used to target *unc-31* (TTTTCAGGAGGATCATGATT) was designed and ordered using the online tool provided by https://sgidtdna.com. Asymmetric hybrid donor repair templates were amplified that contain the GFP-TEV-degron-FLAG coding sequence from the pJW2086 plasmid (Ashley et al., 2021) as previously described (Dokshin et al., 2018). Ultramer primer used: (F: tttcttaaacatttactttttatgttttttttttcttttgcagtcttcgtttggaacatcATGTTAGGAGCAAGTAGTAGTGAAGAAGAAGACGACGATTTTCAGGAGGATCATGATTCTGGTGGCGGTGGATCGGGAGG; R:tgcagaggctctttagagcaaatttttggcagccaacaaaaatacatttcggcttaattttaccacctaattttattctccaaaaacaatatttacactttgaaataaaaaacttcttacCTTGTCATCGTCATCCTTGT). The following mix was then injected into wild-type animals: 4 μg repair template, 4 μg Cas9 protein, 2 μg universal tracrRNA, 1.1 μg crRNA, myo-2::mCherry plasmid (3 ng/μl). F1 progeny were individually isolated and F2 progeny screened for *degron::GFP* knock-in by PCR. After confirmation by Sanger sequencing, the knock-in strain was backcrossed three times to N2 (wild-type) prior to analysis.

### CRISPR-mediated generation the *Pgcy-9::TIR-1* strain

*Pgcy-9-TIR1-F2A-mTagBFP2-C1* vector and strain generation: The *gcy-9* promoter (1918 bp) was amplified from *C. elegans* genomic DNA by PCR using primers: F = AAACGACGGCCAGTGATGCGAGGAAAACTAATGGAGGC R = TTCTCTTTTGCATCGTTATCATCTCATATGAGGGGCATT. Using In-Fusion HD Cloning (Takara Bio), the *gcy-9* promoter was inserted into the pDD357(*Psun-1-TIR1_F2A_mTagBFP2_C1*) vector (kind gift of Jordan Ward) that had been digested with SphI and ClaI. The resultant sequence-confirmed RJP737 *Pgcy-9-TIR1-F2A-mTagBFP2-C1* vector was injected into wild-type *C. elegans* at 25 ng/μl with the following vectors: pCFJ2474(Cas9-expressing vector) at 25 ng/μl (Aljohani et al., 2020), pJW1883(ttTi5605 site targeting sgRNA (F+E) with *R07E5*.*16 U6* promoter and 3’UTR vector) at 25 ng/μl and the *Pmyo-2::RFP* vector at 5 ng/μl. Correct insertion of the *Pgcy-9-TIR1-F2A-mTagBFP2-C1* sequence into the ttTi5605 transposon site on chromosome II was confirmed as described (Ashley et al., 2021). The resultant strain RJP5254 was backcrossed twice with N2 prior to crossing into *rp166*.

### Auxin-inducible degradation (AID) plates

NGM plates were supplemented with 1mM Auxin (indole-3-acetic acid, Alfa Aesar, ALFA10556) from a stock of 400 mM dissolved in ethanol. NGM plates supplemented with ethanol were used as a control.

### Fluorescence microscopy

Animals were anaesthetized with 20 mM NaN_3_ on 5% agarose pads, and images obtained with a Axio Imager M2 fluorescence microscope and Zen software (Zeiss). Fluorescence levels were measured using FIJI software (ImageJ v1.47n).

### Immunoprecipitation and Western Blot

A packed volume (1ml) of *unc-31(rp166); Prgef-1::TIR1* one-day adult hermaphrodites were collected and washed with M9 buffer three times. Worms were pelleted by centrifugation, 3ml of lysis buffer (50 mM Tris pH 7.4, 150 mM NaCl, 2% triton X-100, 1x protease inhibitor cocktail (cOmplete™) was added and solution was sonicated using Bioruptor® (Diagenode). Following centrifugation the supernatant was collected and anti-FLAG antibody (F1804, Sigma-Aldrich) binding Dynabeads™ were used to precipitate GFP-TEV-degron-FLAG-UNC-31. After mixing with 1xBolt™ LDS sample buffer (Invitrogen™) and 1xBolt™ Sample reducing Agent (Invitrogen™), the input sample and post-immunoprecipitation Dynabeads™ were boiled at 95°C for 5 min, then loaded on an 8% polyacrylamide gel for Western Blot. The gel was blotted to a PVDF membrane using the iBlot semi-dry blot system (Thermo Fisher). After blocking with 5% BSA, the PVDF membranes were incubated with anti-FLAG antibody (F1804, Sigma-Aldrich), or anti-actin antibody (MAB1501, Sigma-Aldrich), followed by incubation with horseradish peroxidase-conjugated secondary antibody (61-6520, Thermo Scientific). ECL reagents (Thermo Fisher) and Biorad ChemiDoc XRS+ with CCD camera were used for imaging.

### Locomotion assays

Locomotion speed was measured using the WormLab Imaging System (MBF Bioscience, VT, USA). NGM plates were seeded with 50 μl OP50 bacteria and dried for 1 hr. Six one-day old adults (20 hrs post-mid L4) were picked from auxin or ethanol treatment plates to each assay plate and allowed to acclimate for 10 minutes. Videos were recorded for 1 minute (3.75 frames/second).

Speed = distance travelled/time.

### Egg-laying assay

Young adult hermaphrodites (36 hrs after the mid L4 stage) were individually transferred to a drop of bleach solution (50% commercial bleach/50% 1 M NaOH) on a glass slide. The bleach solution dissolves the hermaphrodite to enable the number of eggs, which are protected by the eggshell, to be counted under DIC optics with a Axio Imager M2 fluorescence microscope.

### Oil-Red O fat staining

*Worm synchronisation:* 5 L4 larvae were picked per genotype to 8 NGM freshly coated with OP50, and incubated for 5 days. Worms were washed from the plates in 4 mL of M9 buffer into a 15 mL conical tube (Thermo Scientific) and eggs were isolated by adding 0.5 mL bleach (White King Premium) + 0.5 mL 5 M NaOH, mixing and incubating for 4 minutes. M9 was added to fill the tube, then samples were centrifuged for 1 minute at 1000 RCF, and the supernatant removed. Egg pellets were washed a further 3 times in M9 buffer, centrifuging for 1 minute at 1000 RCF. The egg pellet was resuspended in ∼1 mL M9 and passed through a 40 µm filter to a fresh 15 mL tube. Samples were kept at room temperature, with gentle rocking for 24 hours to allow all worms to hatch and reach the L1 stage. Approximately 600 L1s were plated onto fresh auxin or ethanol plates (10 × 35mm plates per condition, per genotype) and incubated at 20ºC for 64 hours to reach young adulthood. *Staining:* Oil-Red O stock solution was prepared by adding 0.5 g Oil-Red O with 100 mL isopropanol. The solution covered with foil and mixed at room temperature for 2 days. Immediately prior to commencing fat staining, Oil-Red O stock solution was diluted to 60% with sterile milli-Q filtered water, covered with foil and rotated at room temperature until required. Worms were washed from plates with 10 mL PBS and placed into a 15 mL conical tube using a glass Pasteur pipette and centrifuged at 3300 RCF for 30 seconds. The supernatant was aspirated to 1 mL volume, worms were resuspended and transferred, with Pasteur pipette, to a 1.5 mL microcentrifuge tube. Worms were centrifuged at 3300 RCF for 30 seconds, supernatant removed to 100 µl volume, and pellet resuspended in 1mL PBS with a brief, gentle vortex mix. Samples were transferred to a 1.5ml tube and centrifuged at 3300 RCF for 30 seconds. The supernatant was aspirated to 100 µl, 1 mL PBS was added and then briefly vortexed. Wash steps were repeated for a total of three washes. After removing the final supernatant to 100 µl volume, 1mL of 60 % isopropanol was added to the worm pellet, and samples were incubated for 20 minutes with rotation at room temperature. Samples were centrifuged 3300 RCF for 30 seconds and maximum supernatant was removed without disturbing the worm pellet. Oil-Red O working solution was filtered through a 0.22 µm filter, then 400 µl was added to each worm pellet. Samples were covered in foil, then incubated with rotation overnight at room temperature. The following day, the stained worms were washed twice in PBS + 0.01% Triton-X100, then washed once in PBS, centrifuging at 3300 RCF for 30 seconds. The supernatant was removed, then worms were resuspended and mounted onto agarose pads for imaging. *Quantification:* Oil-Red O staining was quantified in FIJI (ImageJ) by tracing the first 4 intestinal cells proximal to the pharynx. The intensity in these cells was measured in the inverted green channel (where Oil-Red O absorbs the light), collecting area, mean grey value and integrated density data. A region outside of the worm was used as background measurement for Oil-Red O staining and used to calculate the CTCF. CTCF is calculated as: *Integrated Density - (area * mean gray of background)*.

### Electrophysiology recording

NGM plates (60 mm) were seeded with 250 μL OP50 mixed with 5 μL 100 mM all-trans retinal (Sigma) as previously described (Piggott et al., 2011). Seeded retinal plates were stored in the dark at 4°C for up to one week. *C. elegans* hermaphrodites were transferred to retinal plates and grown in the dark for additional 6 hrs before electrophysiological recording. All patch-clamp electrophysiology was performed with dissected adult hermaphrodites under an Olympus microscope (BX51WI) with an EPC-10 amplifier and the patchmaster software (HEKA), using a protocol previously described (Richmond et al., 1999). Each worm was glued on a sylgard-coated coverslip along the dorsal side with a medicalgrade, cyanoacrylate-based glue (Dermabond Topical Skin Adhesive, Ethicon). A small piece of cuticle on the worm body was carefully cut and glued down to the coverslip to expose the BWMs for recording. Current data was filtered at 2 kHz and sampled at 20 kHz. Series resistance and membrane capacitance were both compensated. Recording pipettes were pulled from thick-walled, borosilicate glass (BF150-86-10, Sutter Instruments, USA) and had resistances of 3 - 5 MΩ. The regular bath solution contained (in mM) 140 NaCl, 5 KCl, 5 MgCl_2_, 1 CaCl_2_, 11 glucose, 10 HEPES (330 mOsm, pH adjusted to 7.3). The pipette solution contained (in mM) 120 KCl, 20 KOH, 4 MgCl_2_, 5 EGTA, 0.25 CaCl_2_, 10 HEPES and 5 Na_2_ATP (325 mOsm, pH adjusted to 7.2 using NaOH). BWMs were clamped at −60 mV. Channelrhodopsin was excited with a blue light pulse (3 ms) generated with a blue light-emitting diode (LED) source (Thorlabs, M00552407, Newton, NJ, USA). The light switch was controlled by TTL signals from a HEKA EPC-10 double amplifier. All experiments were performed at room temperature (20 □ - 22 □).

### Statistical analyses

Experiments were performed in three independent replicates with the experimenter blinded to genotype/condition. Statistical analysis was performed in GraphPad Prism 7 using either one-way analysis of variance (ANOVA) when more than two samples were being compared, or Welch’s t-test when two samples were being compared. Values are expressed as mean ±SEM. Differences with a *p* value < 0.05 were considered significant.

## Results and Discussion

### Generation of a conditional degradation allele for UNC-31/CAPS

The *unc-31* gene encodes multiple isoforms. We elected to tag the longest isoforms that contain all the recognized functional domains (Figure 1A-B) (Speese et al., 2007). Expression of N-terminally tagged UNC-31 was previously shown to support transgenic rescue of *unc-31* mutant phenotypes (Speese et al., 2007). Therefore, we used CRISPR-Cas9 engineering to insert a GFP-degron-TEV-3xFLAG cassette at the N-terminus (30 amino acids downstream of the ATG codon) of UNC-31 (Dokshin et al., 2018). The resultant *unc-31(rp166)* knock-in strain was sequenced to confirm the insertion, backcrossed three times with wild-type animals, and re-isolated. Next, we analyzed GFP fluorescence in *unc-31(rp166)* hermaphrodites and *unc-31(rp166)*; *him-5(e1490)* males (Figure 1C). We first detected expression in the nerve ring of 3-fold embryos just prior to hatching (Figure 1C). In adults, strong fluorescence was maintained in the nerve ring and expression was also detected in the ventral and dorsal nerve cords (VNC/DNCs). In males, a similar expression pattern was observed, albeit with additional expression in the male-specific post-cloacal sensilla (Figure 1C). Focusing on *unc-31(rp166)* expression in VNC motor neurons, we observed punctate expression reminiscent of UNC-31 immunostaining representing synapses (Figure S1) (Gracheva et al., 2007). UNC-31 synaptic localization is dependent on the UNC-104 kinesin motor protein (Gracheva et al., 2007). We found that *unc-31(rp166)* expression in VNC motor neurons and the nerve ring is also disrupted in *unc-104(e1265)* mutant animals, with GFP accumulation in cell bodies (Figure S1). Thus, *unc-31(rp166)* expression is detected in the nervous system of both sexes, with the most prominent expression in the nerve ring.

**Figure 1.**
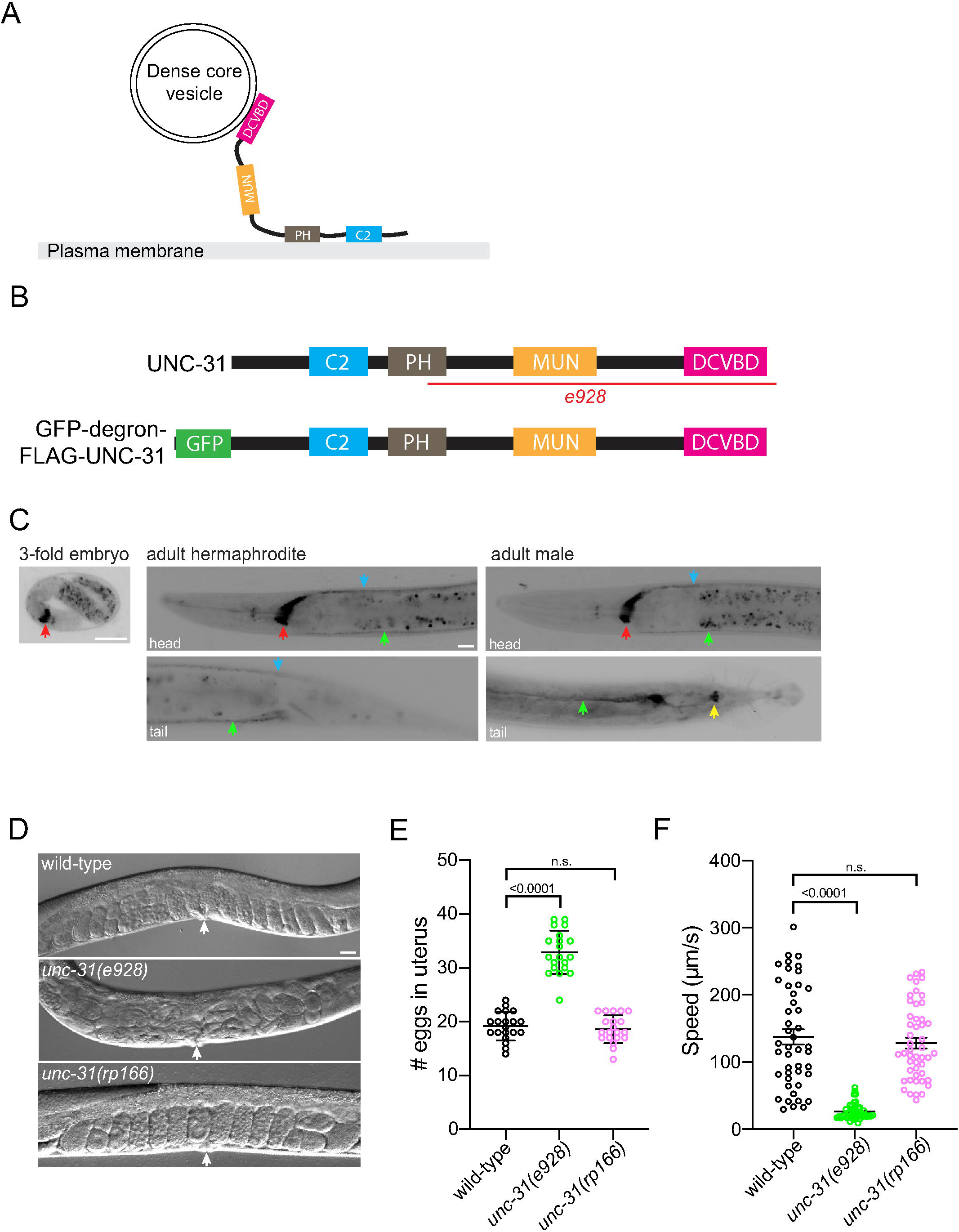
Generation of an UNC-31 conditional degradation allele. (A) Schematic of the UNC-31 protein. C2 = coiled coil domain; PH = pleckstrin homology domain; MUN = Munc13-homology domain; DCVBD = dense-core vesicle binding domain. (B) Schematic of wild-type UNC-31 protein (top) and GFP-degron-FLAG-UNC-31 protein (bottom). *e928 =* deleted region in *unc-31* null mutant (red line) (C) Fluorescent micrographs of *unc-31(rp166)* knock-in GFP expression in a 3-fold embryo (left), adult hermaphrodite (center) and adult male (right). Red arrows = nerve ring; green arrows = ventral nerve cord; blue arrows = dorsal nerve cord; yellow arrow = post-cloacal sensilla. Lateral views, anterior to the left (except male tail image = ventral view). Scale bars = 20 μm. (D-E) DIC images (D) quantification (E) of egg accumulation in wild-type, *unc-31(e928)* and *unc-31(rp166)* adult hermaphrodites (36 hr post-mid L4). White arrows = vulva. n = 20. Data expressed as mean ± SEM and statistical significance was assessed by one-way ANOVA multiple comparison with Dunnett’s multiple comparisons test. n.s. = not significant. Scale bar = 20 μm. (F) Quantification of locomotory speed (distance travelled/time) of wild-type, *unc-31(e928)* and *unc-31(rp166)* adult hermaphrodites (20 hr post-mid L4). n = 45-48. Data expressed as mean ± SEM and statistical significance was assessed by one-way ANOVA multiple comparison with Tukey’s multiple comparisons test. n.s. = not significant.

Fusion of the GFP-degron-TEV-3xFLAG tag to the endogenous UNC-31 protein may have a detrimental effect on UNC-31 function. We did not observe any overt phenotype in *unc-31(rp166)* animals, however, to examine this possibility further, we analyzed two well-studied paradigms of UNC-31 function: egg-laying and locomotion (Avery et al., 1993). We first examined the steady-state accumulation of eggs in the uterus of wild-type, *unc-31(e928)* null mutant and *unc-31(rp166)* one-day adult hermaphrodites. We found that *unc-31(rp166)* animals exhibit wild-type levels of eggs retention (Figure 1D-E). This contrasts to *unc-31(e928)* null mutant animals, which accumulate significantly more eggs than wild-type animals (Figure 1D-E). Next, we examined locomotion of wild-type, *unc-31(e928)* and *unc-31(rp166)* one-day adult hermaphrodites using WormLab automated tracking (MBF Bioscience). We found that the locomotory speed (distance travelled/time) of *unc-31(rp166)* animals is not significantly different to wild type (Figure 1F). In contrast, *unc-31(e928)* null mutant animals exhibit significantly reduced locomotion (Figure 1F). Taken together our data show that the CRISPR-Cas9 generated *unc-31(rp166)* knock-in strain reveals expression in the nervous system and that *in vivo* tagging of UNC-31 does not cause detectable defects in egg-laying or locomotion.

### Temporal degradation of UNC-31

To determine whether UNC-31 can be efficiently depleted by the auxin-inducible degradation system, we introduced a transgene into *unc-31(rp166)* animals that expresses TIR1 in the nervous system under control of the *rgef-1* promoter (Figure 2) (Ashley et al., 2021). We exposed either *unc-31(rp166)* or *unc-31(rp166); Prgef-1::TIR1* animals to auxin for 5 hr and measured GFP fluorescence in the nerve ring (Figure 2A-B). We found that GFP levels were unchanged in *unc-31(rp166)* animals exposed to auxin compared to the ethanol control (Figure 2A). In contrast, auxin robustly depleted GFP fluorescence in *unc-31(rp166); Prgef-1::TIR1* animals (Figure 2B). We confirmed auxin-induced depletion of *unc-31(rp166); Prgef-1::TIR1* animals by probing for the FLAG tag by Western blot, finding that GFP-degron-FLAG-UNC-31 protein was undetectable after 24 hr of auxin treatment (Figure 2C).

**Figure 2.**
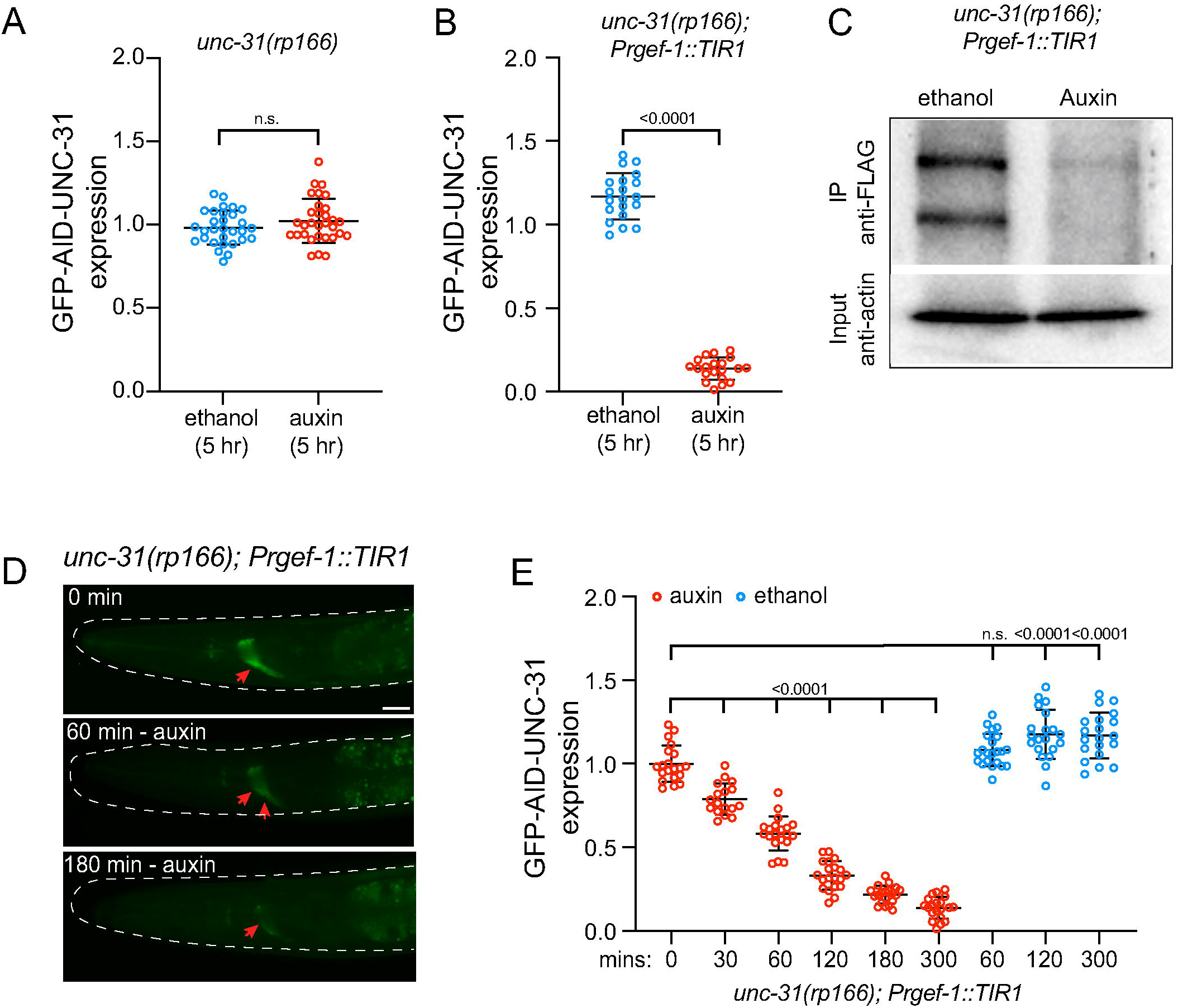
Temporal knockdown of UNC-31 expression with auxin. (A-B) Quantification of GFP expression in the nerve ring of *unc-31(rp166)* (A) and *unc-31(rp166); Prgef::TIR1* (B) mid-L4 larvae exposed to 1 mM auxin or ethanol for 5 hr, relative to untreated controls. n = 30-31 in A, 20 in B. Data expressed as mean ± SEM and statistical significance was assessed by unpaired t-test. n.s. = not significant. (C) Western blot of immunoprecipitated GFP-degron-FLAG-UNC-31 protein from *unc-31(rp166); Prgef::TIR1* animals exposed to ethanol or auxin for 24 hr. Two major UNC-31 bands were detected. (D) Fluorescent micrographs of GFP expression in *unc-31(rp166); Prgef::TIR1* L4 hermaphrodites following 1 mM auxin exposure for 0, 60 and 180 min. Red arrows = nerve ring; dotted line = worm outline. Scale bar = 20 μm. (E) Quantification of GFP expression in the nerve ring of *unc-31(rp166); Prgef::TIR1* in mid-L4 larvae exposed to 1 mM auxin or ethanol for the specified time periods in minutes. n = 17-21. Data expressed as mean ± SEM and statistical significance was assessed by one-way ANOVA multiple comparison with Tukey’s multiple comparisons test. n.s. = not significant.

The successful utilization of *unc-31(rp166); Prgef-1::TIR1* animals in behavioral experiments may depend on the rapidity of depletion. We therefore analyzed the kinetics of auxin-mediated depletion of UNC-31 over a 5-hr period by measuring nerve ring fluorescence in L4 larvae (Figure 2D-E). We found that within 30 min of auxin exposure UNC-31 expression reduced by ∼20% (Figure 2E) and observed gradual reduction of UNC-31 expression over time to ∼90% of control levels after 5 hr of auxin exposure (Figure 2D-E). We found that ethanol, the diluent for auxin that is used as a control, caused a slight but significant increase in UNC-31 expression after 1 hr (Figure 2E). This should be considered when performing experiments with this strain.

### Conditional depletion of UNC-31 causes defects in egg-laying and locomotion

To determine the utility of the *unc-31(rp166)* tool for functional studies, we assayed egg-laying and locomotion (Figures 3 and 4). We exposed wild-type, *unc-31(rp166)* and *unc-31(rp166); Prgef-1::TIR1* mid-L4 stage hermaphrodites to auxin or ethanol for 24 hr and counted egg retention in the uterus (Figure 3). We found that the accumulation of eggs in wild-type and *unc-31(e928)* animals is not significantly affected by auxin (Figure 3B). In contrast, auxin caused significant accumulation of eggs in *unc-31(rp166); Prgef-1::TIR1* animals (Figure 3A-B).

**Figure 3.**
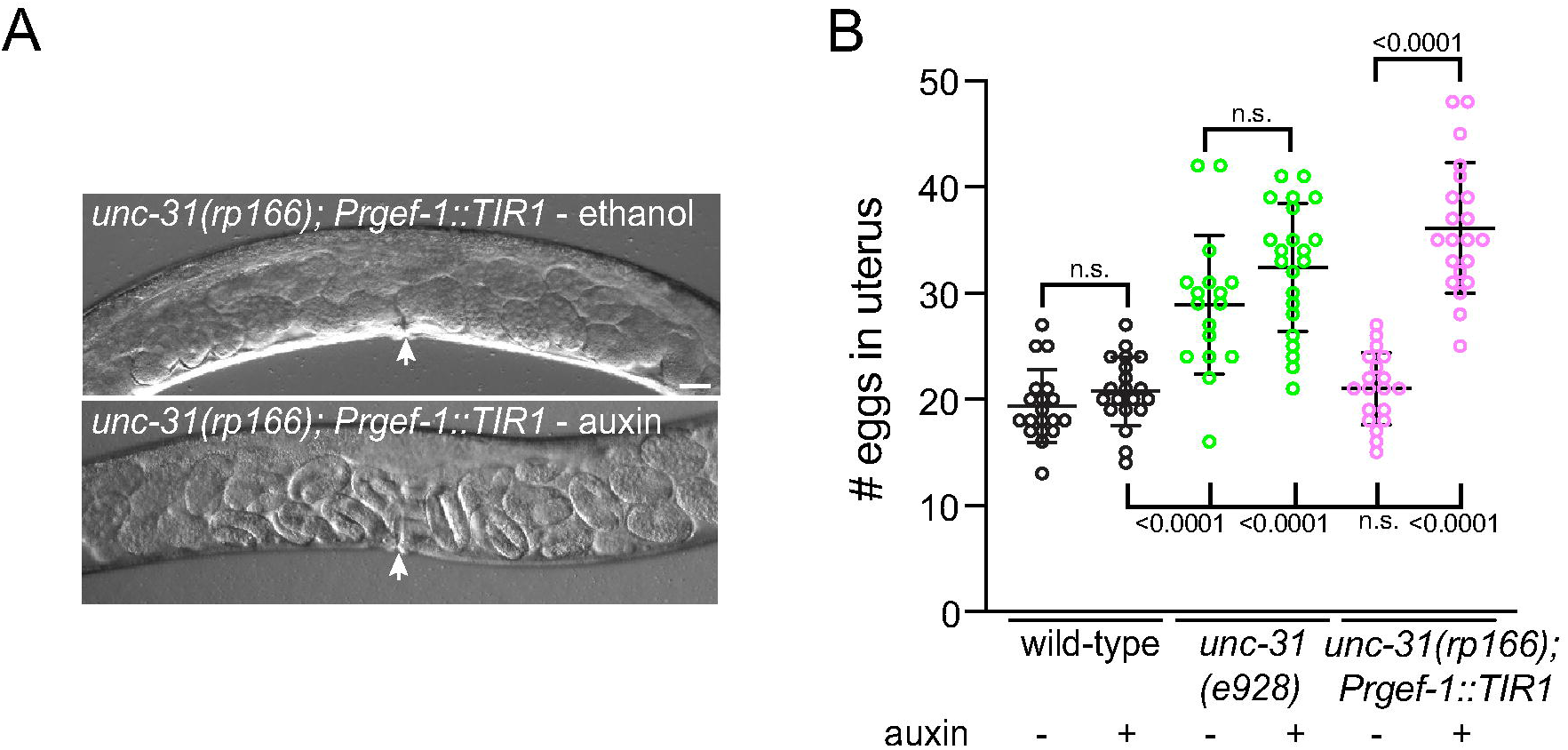
Auxin-induced UNC-31 knockdown causes egg accumulation. (A) DIC images of *unc-31(rp166); Prgef::TIR1* 1-day adult hermaphrodites exposed to ethanol or auxin for 24 hr from the mid-L4 larval stage. White arrows = vulva. Scale bar = 20 μm. (B) Quantification of egg accumulation in wild-type, *unc-31(e928)* and *unc-31(rp166); Prgef::TIR1* 1-day adult hermaphrodites. Auxin and ethanol treatments were for 24 hr from the mid-L4 larval stage. n = 17-22. Data expressed as mean ± SEM and statistical significance was assessed by one-way ANOVA multiple comparison with Dunnett’s multiple comparisons test. n.s. = not significant.

**Figure 4.**
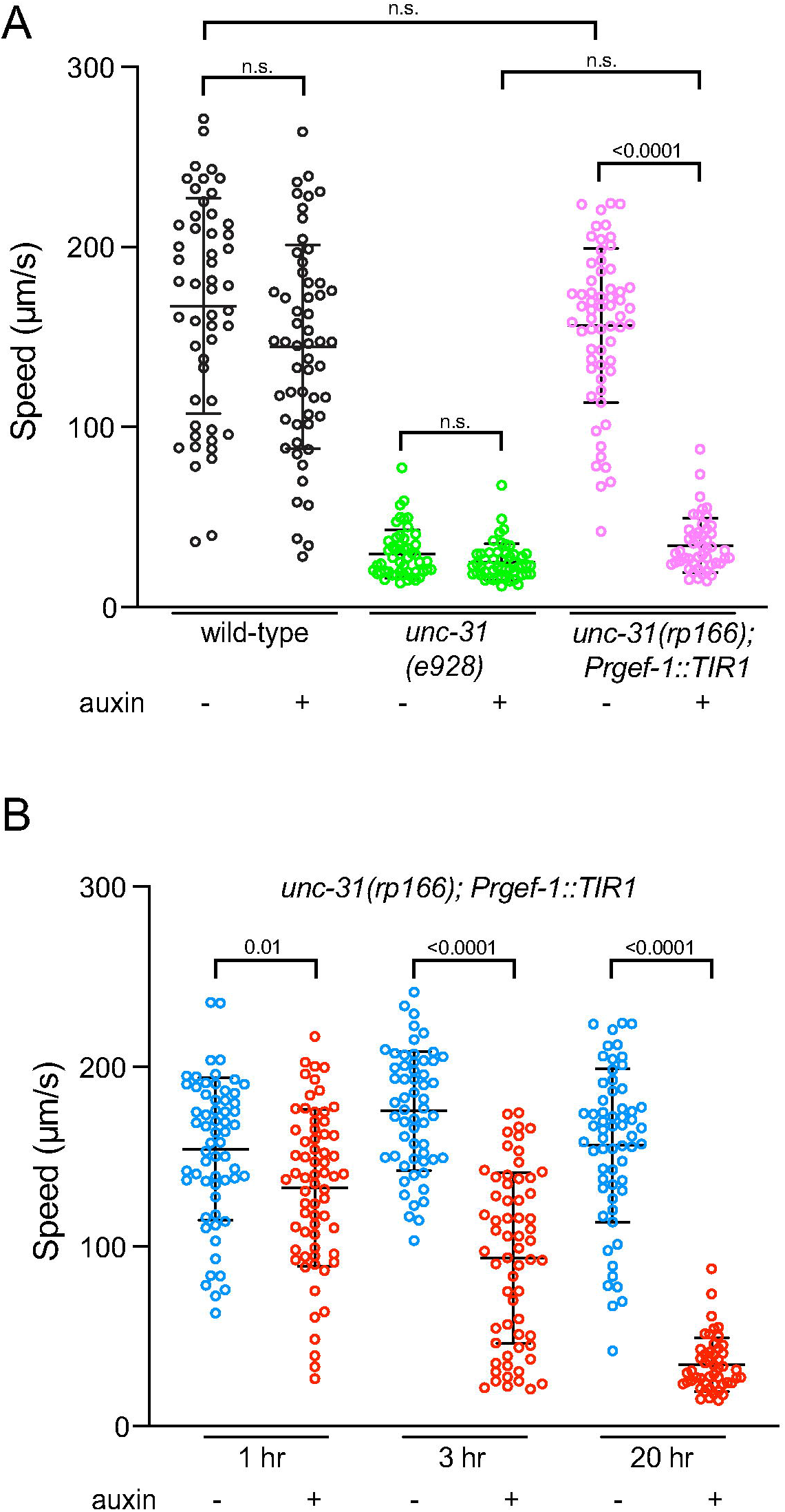
Auxin-induced UNC-31 knockdown causes defects in locomotion. (A) Quantification of locomotory speed (distance travelled/time) of wild-type, *unc-31(e928)* and *unc-31(rp166); Prgef::TIR1* adult hermaphrodites exposed to 1 mM auxin or ethanol for 20 hr from the mid-L4 stage. n = >50. Data expressed as mean ± SEM and statistical significance was assessed by one-way ANOVA multiple comparison with Sidak’s multiple comparisons test. n.s. = not significant. (B) Quantification of locomotory speed (distance travelled/time) of *unc-31(rp166); Prgef::TIR1* adult hermaphrodites exposed to 1 mM auxin or ethanol for 1 hr, 3 hr or 20 hr. n = >50. Data expressed as mean ± SEM and statistical significance was assessed by one-way ANOVA multiple comparison with Sidak’s multiple comparisons test. n.s. = not significant.

Next, we examined locomotion of wild-type, *unc-31(e928)* and *unc-31(rp166); Prgef-1::TIR1* one-day adult hermaphrodites that were exposed to auxin for 20 hr from the mid-L4 stage (Figure 4). Using WormLab automated tracking (MBF Bioscience) we found that the locomotory speed of wild-type and *unc-31(e928)* animals was not affected by auxin exposure (Figure 4A). In contrast, *unc-31(rp166); Prgef-1::TIR1* exposed to auxin exhibited locomotion similar to the *unc-31(e928)* null mutant (Figure 4A). This shows that UNC-31 is continually required for *C. elegans* locomotion. To examine the kinetics of locomotory decline following auxin-induced UNC-31 depletion, we measured the locomotion speed of *unc-31(rp166); Prgef-1::TIR1* adult hermaphrodites exposed to auxin for 1 hr, 3 hr and 20 hr (Figure 4B). We found that within 1 hr of auxin exposure, when ∼40% of UNC-31 protein is depleted (Figure 2E), there is a significant decrease in locomotion speed compared to control animals (Figure 4B). After 3 hr of auxin exposure the locomotory speed of *unc-31(rp166); Prgef-1::TIR1* animals reduces by ∼60%, with a further decrease to levels of *unc-31* null mutant locomotion within 20 hr (Figure 4A-B). Together, these data show that *unc-31(rp166)* enables temporal depletion of UNC-31 protein to generate robust behavioral phenotypes.

### Conditional depletion of UNC-31 causes reduced channelrhodopsin-induced currents at the neuromuscular junction

UNC-31 is a known regulator of DCV release (Gracheva et al., 2007; Sieburth et al., 2007). However, it was previously shown that *unc-31* null mutants also exhibit a smaller evoked response to electrical stimulation (Gracheva et al., 2007) and blue light-induced evoked currents in animals expressing channelrhodopsin (Lindsay et al., 2011). To examine whether auxin-induced depletion of UNC-31 causes defects in vesicle release, we performed electrophysiological recording at the cholinergic neuromuscular junctions (NMJs) (Liu et al., 2018). We expressed channelrhodopsin in cholinergic motor neurons (*oxIs364 [Punc-17::ChR2]*) and examined evoked currents following blue-light stimulation (Liewald et al., 2008; Liu et al., 2009). As previously reported, we found that *unc-31(e928)* null mutant animals expressing channelrhodopsin in cholinergic motor neurons exhibit smaller evoked currents compared to wild-type animals (Figure 5A-C) (Gracheva et al., 2007). Next, we examined blue light-induced inward currents in *unc-31(rp166); Prgef-1::TIR1; oxIs364* animals exposed to auxin or the ethanol control for 6 hrs. We found that *unc-31(rp166); Prgef-1::TIR1; oxIs364* animals exposed to ethanol exhibit a similar response as wild type animals (Figure 5). In contrast, *unc-31(rp166); Prgef-1::TIR1; oxIs364* animals exposed to auxin show a significant decrease in evoked amplitudes that is comparable to *unc-31(e928)* null mutant animals (Figure 5). These data show that auxin-induced depletion of UNC-31 causes defects in vesicle release.

**Figure 5.**
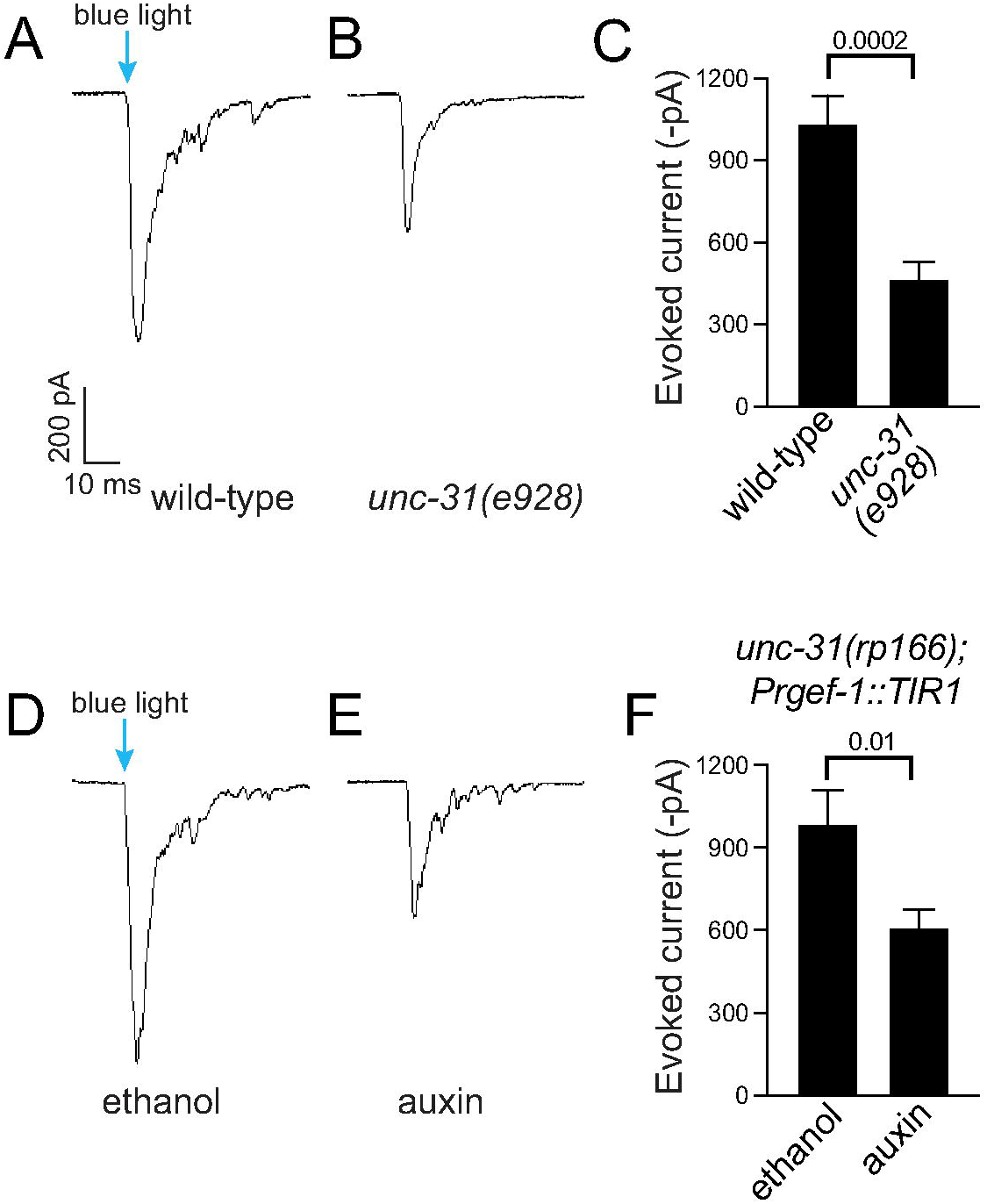
Auxin-induced UNC-31 knockdown reduces channelrhodopsin-induced currents at the neuromuscular junction. (A-C) Sample traces (A-B) and quantification (C) of blue light stimulated amplitude of evoked EPSCs of wild-type (A) and *unc-31(e928)* (B) adult hermaphrodites. n = 9-11. (D-F) Sample traces (D-E) and quantification (F) of blue light stimulated amplitude of evoked EPSCs of ethanol (D) and auxin (E) treated *unc-31(rp166); Prgef::TIR1* adult hermaphrodites. n = 6-8. Data expressed as mean ± SEM and statistical significance was assessed by unpaired t-test.

### BAG neuron-specific depletion of UNC-31 causes intestinal fat accumulation

Next, we examined the utility and spatial resolution of the UNC-31 conditional depletion tool. The BAG sensory neurons are key regulators of *C. elegans* behavior and physiology (Zimmer et al., 2009; Brandt et al., 2012; Carrillo et al., 2013; Juozaityte et al., 2017; Handley et al., 2022). We previously showed that animals lacking the BAG neuron fate-determining transcription factor ETS-5 store excess intestinal fat (Juozaityte et al., 2017). In addition, BAG-specific *unc-31* RNAi or loss of the INS-1 peptide from the BAGs results in increased fat storage (Juozaityte et al., 2017; Handley et al., 2022). To examine whether the conditional auxin system could be used to confer neuron-specific phenotypes, we expressed TIR1 specifically in the BAG neurons using the *gcy-9* promoter (Brandt et al., 2012) and crossed this strain into *unc-31(rp166)*. We exposed wild-type and *unc-31(rp166); Pgcy-9::TIR1* animals to ethanol or auxin from L1 larvae to adult and measured fat levels by Oil-Red O (ORO) staining (Figure 6). We found that BAG-specific UNC-31 depletion increases intestinal fat storage levels (Figure 6). These data confirm the overall inhibitory effect of BAG-derived neuropeptides on intestinal fat levels.

**Figure 6.**
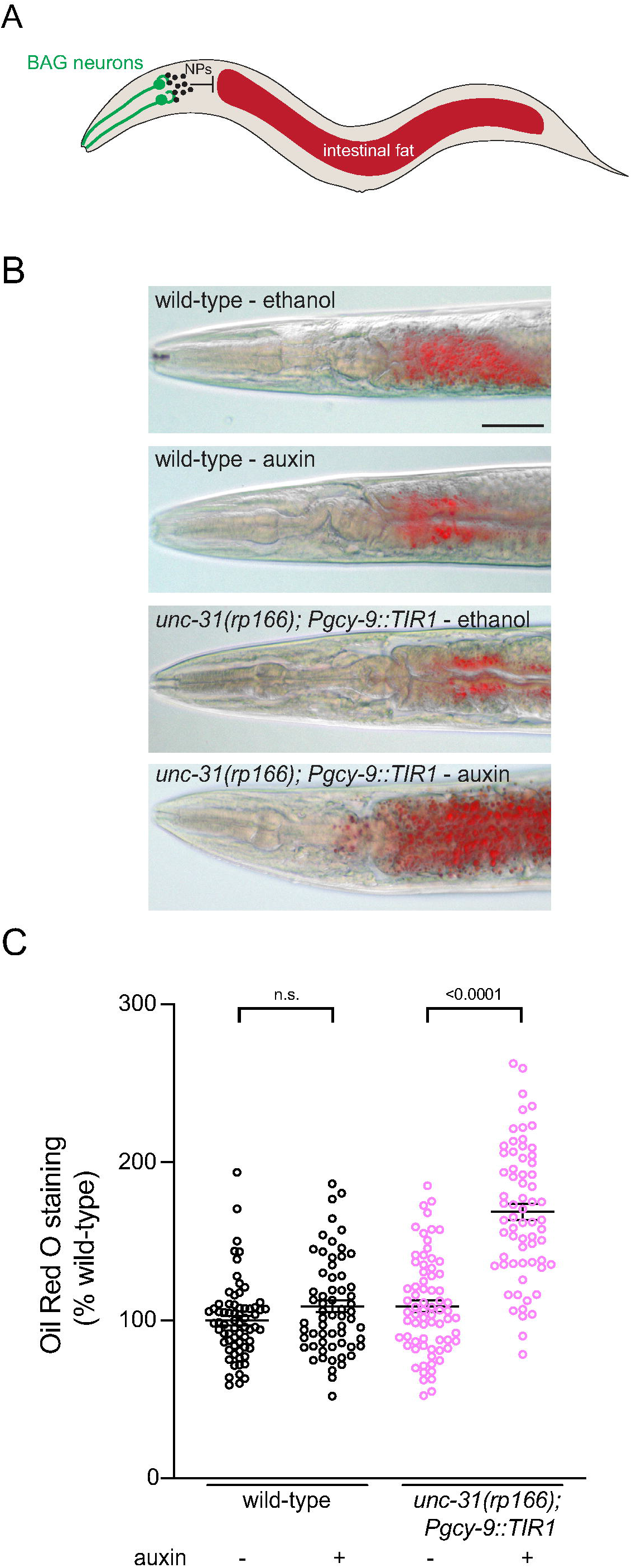
BAG neuron-specific UNC-31 knockdown increases intestinal fat levels. (A) Schematic of *C. elegans* showing that neuropeptides (NPs) released from the BAG neurons inhibit intestinal fat storage (Juozaityte et al., 2017; Handley et al., 2022). (B-C) Representative images (B) and quantification (C) of Oil-Red O (ORO) staining in wild-type and *unc-31(rp166); Pgcy-9::TIR1* adult hermaphrodites exposed to 1 mM auxin or ethanol from L1 larvae to adult. n = >62. Data presented as ORO intensity as % wild-type, mean ± SEM and statistical significance was assessed by one-way ANOVA multiple comparison with Tukey’s multiple comparisons test. n.s. = not significant. Scale bar 50 μm.

In conclusion, we describe here the generation and validation of a genetic tool that enables the function of the UNC-31/CAPS protein to be dissected in a spatiotemporal manner. In combination with neuron-specific expression of the *A. thaliana* TIR1 protein, this tool will enable investigators to inhibit neuropeptide release in a highly controlled manner. This will facilitate functional studies of individual post-mitotic neurons, which will help to determine the roles of individual neurons and circuits in controlling behavior and physiology.

## Acknowledgements

We thank members of the Pocock laboratory for comments on the manuscript. We thank Jordan Ward for providing the pPD357 vector prior to publication. Some strains were provided by the Caenorhabditis Genetics Center (University of Minnesota), which is funded by NIH Office of Research Infrastructure Programs (P40 OD010440). This work was supported by the following grants: NHMRC (GNT1105374 to RP) and a veski Innovation Fellowship (VIF23 to RP).

## Supplementary Information

**Figure S1.**
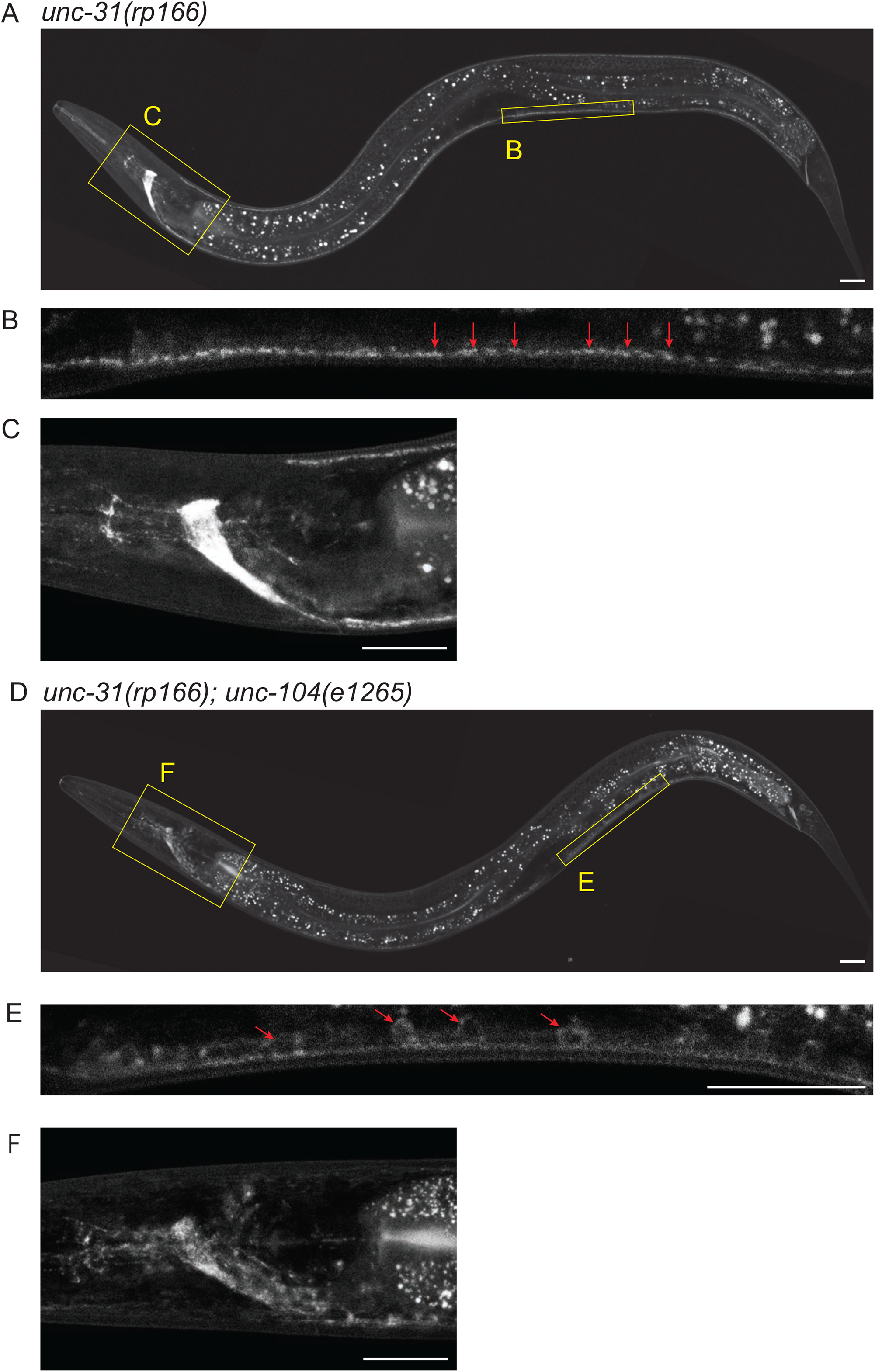
Localization of endogenously-tagged UNC-31 is dependent on the kinesin UNC-104. (A-F) Fluorescent confocal images of *unc-31(rp166)* knock-in GFP expression in wild-type (A-C) and *unc-104(e1265)* (D-F) adult hermaphrodites. Yellow boxes in (A and D) expanded in (B-C and E-F). Arrows show punctate localization (B) compared to cell body accumulation (E). Lateral views, anterior to the left. Scale bars = 20 μm.

**Table S1. Strains used in this study**

## Notes

**Conflicts of interest:** The authors declare no competing financial interests

### Competing Interest Statement

The authors have declared no competing interest.

